# LEM-3 is a midbody-tethered DNA nuclease that resolves chromatin bridges during cytokinesis

**DOI:** 10.1101/221291

**Authors:** Ye Hong, Remi Sonneville, Bin Wang, Viktor Scheidt, Bettina Meier, Alexander Woglar, Sarah Demetriou, Karim Labib, Verena Jantsch, Anton Gartner

**Author notes:** Current address, Departments of Developmental Biology and Genetics, Stanford University School of Medicine, Stanford, USA.

## Abstract

Faithful chromosome segregation and genome maintenance requires the removal of all DNA bridges that physically link chromosomes before cells divide. Using *C. elegans* embryos we show that the LEM-3/Ankle1 nuclease defines a new genome integrity mechanism by processing DNA bridges right before cells divide. LEM-3 acts at the midbody, the structure where abscission occurs at the end of cytokinesis. LEM-3 localization depends on factors needed for midbody assembly, and LEM-3 accumulation is increased and prolonged when chromatin bridges are trapped at the cleavage plane. LEM-3 locally processes chromatin bridges that arise from incomplete DNA replication, unresolved recombination intermediates or the perturbance of chromosome structure. Proper LEM-3 midbody localization and function is regulated by AIR-2/Aurora B kinase. Strikingly, LEM-3 act cooperatively with the BRC-1/BRCA1 homologous recombination factor to promote genome integrity. These findings provide a molecular basis for the suspected role of the LEM-3 orthologue Ankle1 in human breast cancer.

## Introduction

Faithful chromosome segregation requires all connections that physically link chromosomes to be removed before cells divide; failure to do so may lead to the severing of chromosomes, prevent cell division or cause aneuploidy and polyploidization^1–4^. It is widely known that proteinaceous connections between chromosomes mediated by cohesins must be removed prior to segregation, but so must DNA structures that link chromatids. These DNA connections include DNA repair intermediates, points at which chromatids have become intertwined, and loci that have not fully replicated. Cytologically, connections between separating chromatids can take the form of chromatin bridges or ultrafine DNA bridges^5,6^. The majority of DNA linkages, such as branched recombination intermediates, are ‘dissolved’ by the combined action of the BLM helicase and Topoisomerase III concomitant with DNA replication, or by the MUS81-EME1 nuclease in G2/M and the GEN1 Holliday junction resolvase during anaphase (for review^7^). However, remaining chromatin bridges can persist into telophase. During cytokinesis the Aurora B kinase dependent NoCut checkpoint detects chromatin bridges and delays abscission of daughter cells in order to allow for chromatin bridge resolution^4,8^. This delay is achieved through continuous Aurora B activation, while cytokinesis is completed upon kinase inactivation. However, little is known about how chromatin bridges are resolved when the Aurora B kinase delays cytokinesis.

The LEM-3 nuclease was discovered in a genetic screen for DNA repair genes in *C. elegans*^9^. *lem-3* mutants are hypersensitive to ionizing radiation (IR), UV light and DNA cross-linking agents^9^. However, previous studies have not shown how LEM-3 mediates genome stability. The mammalian orthologue of LEM-3 is known as Ankle1 and is predominantly expressed in hematopoietic tissues. Ankle1-deficient mice are viable and show no detectable defects^10–12^ Interestingly, non-coding polymorphisms in either the ANKLE1 locus or the neighbouring ABHD8 locus have been associated with increased risk of breast and ovarian cancer in the general population and in carriers of BRCA1 mutations, suggesting that altered expression of ANKLE1 or ABHD8 might contribute to tumourigenesis^13–15^.

Here we identify a novel mechanism conferred by LEM-3 nuclease to preserve genome integrity during the final stages of cell division in *C. elegans*. Our data indicates that LEM-3 can be regulated by the Aurora B kinase AIR-2 and provides a final safeguard to resolve the remaining chromatin bridges at the very end of cytokinesis, by virtue of the recruitment of LEM-3 to the midbody.

## Results

### LEM-3 acts outside of known DNA repair pathways

*C. elegans lem-3* null mutants were reported to proliferate normally but are hypersensitive to a range of DNA damaging agents including ionizing irradiation (IR) which predominantly acts by inducing DNA double-strand breaks^9^. The mammalian orthologue of LEM-3, known as Ankle1, is poorly characterised^10–12^. We wished to determine whether LEM-3 acts in previously characterised DNA damage repair pathways, or else defines a new mechanism. We found that after IR treatment, mutation of *lem-3* showed more severe phenotypes when combined with null alleles defective in the three major DNA double-strand break repair modalities: BRC-1 dependent homologous recombination (HR), polymerase Theta (POL-Q) mediated end-joining^16^ and LIG-4 dependent non-homologous end-joining^17^ (Fig. S1A-S1C). Taken together, our genetic data indicate that LEM-3 might act in a previously unknown response mechanism.

### LEM-3 accumulates at the midpoint of chromatin bridges during cell division

To better understand how LEM-3 helps to maintain genome stability, we assessed LEM-3 localization using an existing transgene carrying a *lem-3::YFP* translational fusion, as well as a GFP tagged *lem-3* derivate, generated by CRISPR-Cas9 mediated genome editing. Both constructs complement the DNA damage hypersensitivity phenotypes conferred by *lem-3* null alleles (Fig. 5C)^9^. Consistent with previous findings^9^, LEM-3 appeared excluded from the nucleus, and formed prominent foci of uncertain nature outside the nuclei in the absence of DNA damage (Fig. 1A). To explore potential links between these LEM-3 foci and chromosome segregation, we exposed worms to a range of treatments that induce chromatin bridges during cell division. As shown in Fig. 1B, LEM-3 localized to the center of chromatin bridges that formed between dividing nuclei in response to DNA breaks (IR), DNA replication defects (hydroxyurea treatment or RNAi, targeting the MCM-7 helicase subunit), condensation defects (RNAi, to deplete the HCP-6 condensin I subunit), or decatenation defects (RNAi depletion of the type II topoisomerase TOP-2^18,19^). Careful examination of LEM-3 localization by OMX super-resolution microscopy of DAPI stained embryos treated with *mcm-7* RNAi revealed that LEM-3 initially formed a ring structure encircling chromatin bridges between two separated daughter cells (Fig. 1C, 4A and S7D). In telophase nuclei this structure shrinks accompanied by the gradual elongation of chromatin bridges. Finally, LEM-3 concentrates in the centre of chromatin bridges between late telophase nuclei (Fig. 1C and S7D).

**Fig 1.**
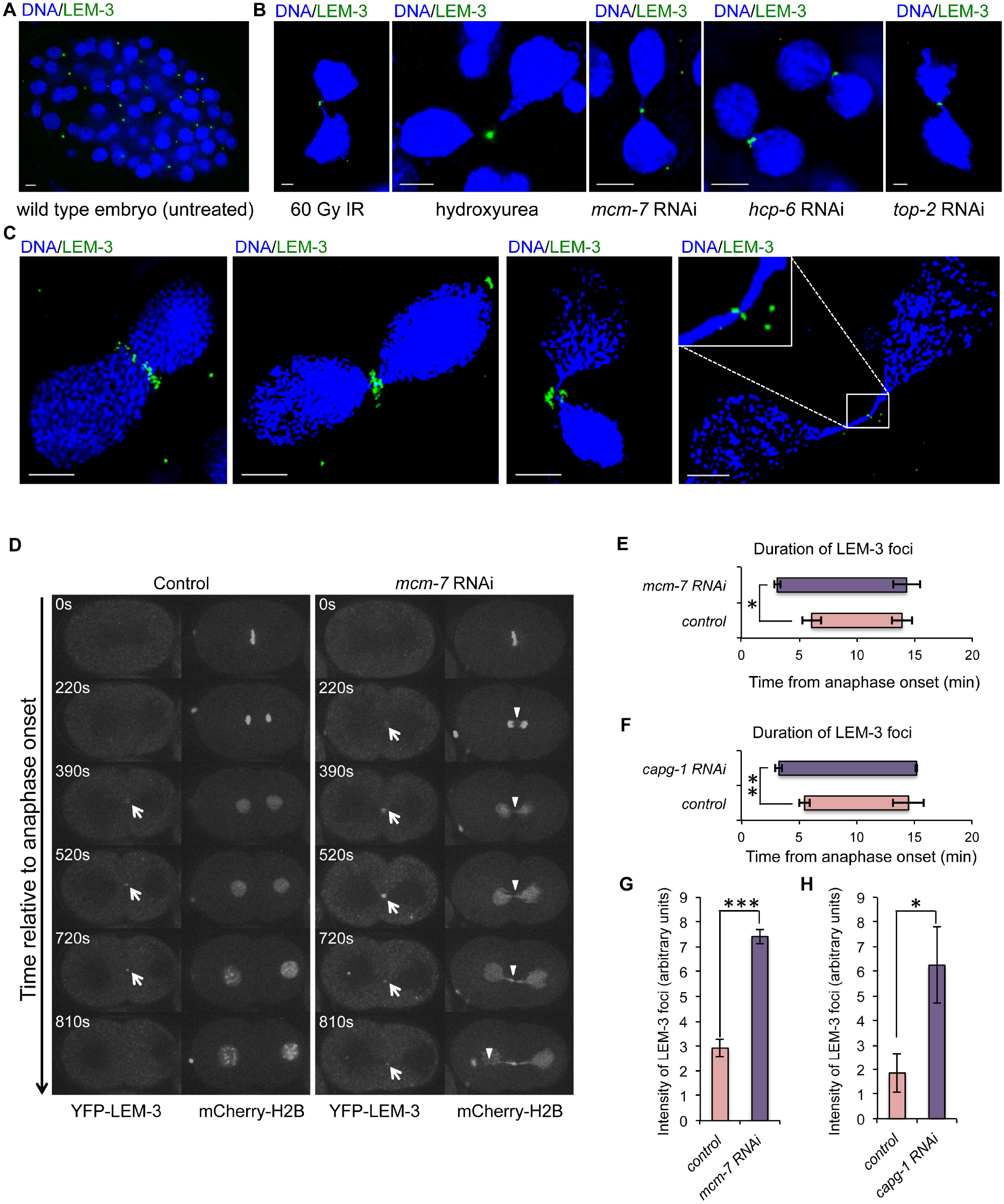
Localization of LEM-3 on chromatin bridges and at the midbody. (A) Localization of YFP-LEM-3 in wild type embryo. (B) Localization of YFP-LEM-3 on chromatin bridges generated by IR, hydroxyurea and RNAi depletion of replicative helicase subunit MCM-7, condensin II component HCP-6 and topoisomerase TOP-2. (C) Representative OMX images of YFP-LEM-3 localization at different stage of cytokinesis in the presence of chromatin bridges induced by *mcm-7* RNAi. Scale bars: 2 μ m. (D) Localization of LEM-3 on chromatin bridges induced by *mcm-7* RNAi. Arrows indicate YFP-LEM-3. Arrowheads indicate the chromatin bridges. Times are relative to anaphase onset of the first division. (E-F) Quantification of the average time duration of YFP-LEM-3 localization at the midzone/midbody during the first mitotic cell division in wild type, *mcm-7* RNAi (E) and *capg-1* RNAi (F) embryos. Error bars represent standard deviation of the mean. (G-H) Quantification of the intensity of YFP-LEM-3 foci during the first mitotic division in wild type, *mcm-7* RNAi (G) (measured at 2.5 min after first appearance of LEM-3 foci) and *capg-1* RNAi (H) (measured at 2 min after first appearance of LEM-3 foci) embryos. Asterisks indicate statistical significance as determined by two-tailed Student T-test. P values below 0.05 were consider significant, where p<0.05 was indicated with *, p<0.01 with ** and p<0.001 with ***.

### LEM-3 is a midbody-tethered protein

Given the LEM-3 staining pattern, we speculated that LEM-3 might localize at the midbody, which defines the site of cell abscission, the complete separation of two daughter cells at the end of cytokinesis. Using a strain carrying YFP-LEM-3 and mCherry-histone H2B fusions we found that YFP-LEM-3 localizes to the midbody region starting from telophase (Fig. 1D, white arrows; Movie S1). To establish if LEM-3 indeed colocalizes with the midbody, we compared the relative localization of LEM-3 and ZEN-4. ZEN-4 is a component of the centralspindlin complex, comprised of the ZEN-4/MKLP1 kinesin 6 motor protein and the CYK-4 Rho family GAP^20^, both essential for midbody formation and cytokinesis. We observed that LEM-3 co-localizes with ZEN-4 at the midbody (Fig. 2A; Movie S2). Furthermore, midbody localisation of LEM-3 could not be observed upon RNAi depletion of CYK-4 or SPD-1, proteins essential for the integrity of the midbody, (Fig. 2B and 2C), even in the presence of chromatin bridges induced by *mcm-7* RNAi (Fig. 2B and 2C). We also investigated if LEM-3 localization required cleavage furrow ingression. As expected, depletion of a contractile ring component NMY-2 inhibited furrow formation and ingression but didn’t affect the central spindle formation (Fig. S3A). We found that LEM-3 could still be detected at the midzone upon *nmy-2* RNAi (Fig. S3A). In summary, the LEM-3 nuclease associates with the midbody, and LEM-3 midbody location depends on the formation of the central spindle and the midbody.

**Fig 2.**
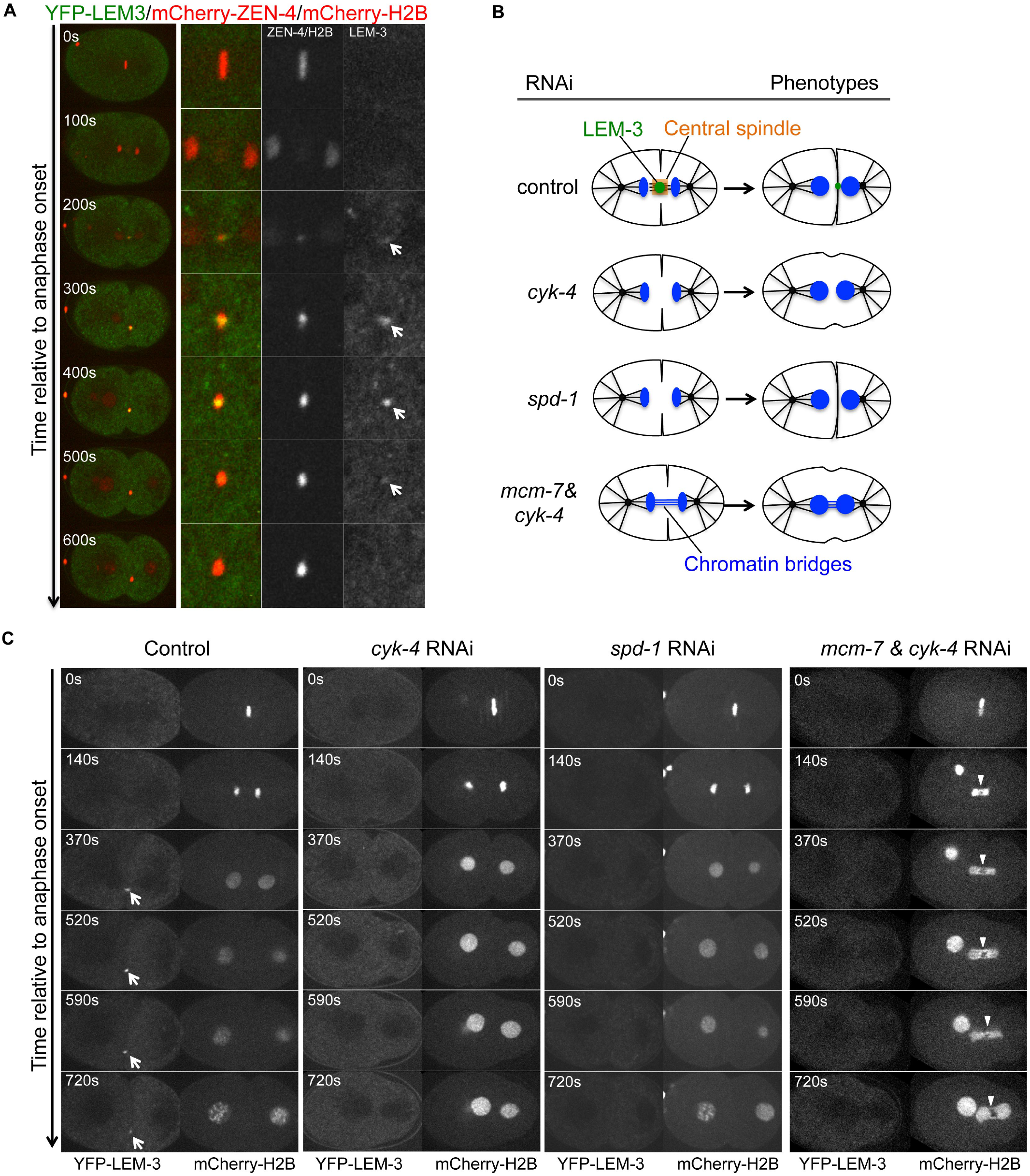
Central spindle formation is required for midbody localization of LEM-3. (A) LEM-3 colocalized with the central spindle component ZEN-4. Arrows indicate YFP-LEM-3. (B) Schematic of LEM-3 localization in control embryos (normal central spindle formation) and *cyk-4, spd-1* RNAi embyros (defective central spindle formation). (C) Midbody localization of LEM-3 depends on the central spindle component CYK-4 and the microtubule-bundling protein SPD-1. Depletion of the central spindle by *cyk-4* RNAi, no LEM-3 foci are detectable even in the presence of chromatin bridges generated by *mcm-7* RNAi. Images were taken from a time-lapse recording of the *cyk-4* RNAi, *spd-1* RNAi, and *cyk-4* combined with *mcm-7* RNAi treated embryos expressing YFP-LEM-3 and mCherry-H2B from the anaphase onset of the first mitotic division. Arrowheads indicate chromatin bridges. Times are relative to anaphase onset.

### LEM-3 is required for the resolution of chromatin bridges

We next sought to determine whether LEM-3 is required to resolve chromatin bridges that result from incomplete DNA replication, defective chromosome condensation, or unresolved recombination intermediates. We found that LEM-3 started to accumulate earlier at the midbody, and the intensity of LEM-3 foci was stronger in the presence of chromatin bridges generated by *mcm-7* RNAi or the condensin I subunit *capg-1* RNAi (Fig. 1D-H, white arrows for LEM-3, white arrowheads for chromatin bridges; Movie S3).

When DNA replication was mildly impaired by weak depletion of the MCM-7 helicase subunit via low dose RNAi (10% bacteria expressing *mcm-7* RNAi), cell cycle progression occurred normally, and no chromatin bridges were observed (Fig. 3A; 10/10 embryos), consistent with our previous work^18^. In contrast, chromatin bridges formed and persisted in all cases (12/12 embryos; Fig. 3A) in *lem-3* mutants. In the absence of replication stress inflicted by *mcm-7* depletion no chromatin bridges formed in *lem-3* mutants (Figure S4A and S5C; 10/10 embryos). It thus appears that DNA bridges arising from incomplete DNA replication are efficiently processed in wild type, but remain unresolved in *lem-3* mutants. Consistent with this view, we found that chromatin bridges were eventually resolved at the end of the first cell division in 33% of wild type embryos exposed to the 100% *mcm-7* RNAi (n=9), while bridges persisted in all *lem-3* mutant embryos treated with 100% *mcm-7* RNAi (n=14) (Fig. 3B). Similarly, whereas the chromatin bridges induced by depletion of the Condensin I subunit CAPG-1 were eventually resolved in wild type (n=5), all *lem-3* embryos tested showed persistent unresolved bridges (n=5) (Fig S2A, white arrowheads).

**Fig. 3.**
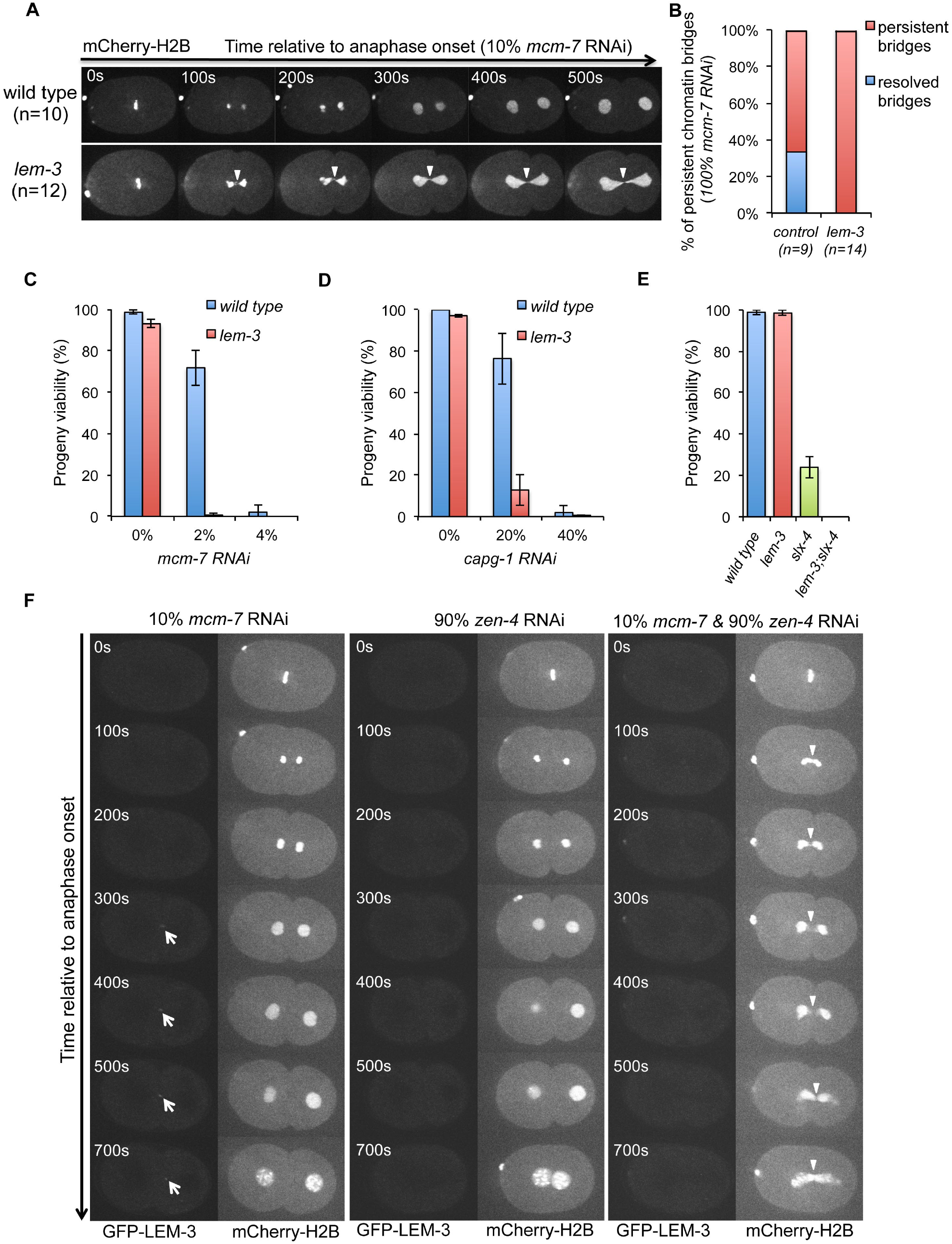
LEM-3 contributes to chromatin bridge resolution. (A) Chromosome segregation in wild type and *lem-3* mutant embryos upon partial depletion of DNA replication helicase subunit MCM-7. Images were taken from embryos expressing mCherry-H2B fed on 10% *mcm-7* RNAi producing bacteria. Arrowheads indicate chromatin bridges. Times are relative to anaphase onset of the first division. (B) Quantification of persistent chromatin bridges in wild type and *lem-3* mutant embryos upon full depletion of MCM-7 (100% *mcm-7* RNAi). n sample size. (CD) Progeny viability of *lem-3* mutants upon partial depletion of MCM-7 (C) and CAPG-1 (D). (E) Genetic interaction between LEM-3 and SLX-4. Progeny viability in % was determined by counting number of viable eggs/total number of eggs laid. Error bars represent standard deviation of the mean. (F) Formation of chromatin bridges upon partial depletion of DNA replication helicase subunit MCM-7 and central spindle component ZEN-4. Arrowheads indicate chromatin bridges.

To further explore the functional significance of chromatin bridge resolution by LEM-3, we examined embryonic viability in either *lem-3* mutants alone or in combination with non-lethal low-doses RNAi against *mcm-7* or *capg-1*. Whereas 72% and 76% of wild type worms survived partial RNAi depletion with *mcm-7* RNAi (2% bacteria expressing *mcm-7* RNAi) or *capg-1* RNAi (20% bacteria expressing *capg-1* RNAi) respectively, the same RNAi treatment led to only 0.6% and 12.8% viable embryos in *lem-3* mutants (Fig. 3C and 3D). The hypersensitivity of *lem-3* mutants to the partial inhibition of MCM-7 and CPAG-1 indicates that LEM-3 is important to process DNA intermediates that result from incomplete DNA replication or partial chromosome decondensation to ensure viability. We then investigated whether the same was true for DNA bridges that result from incomplete processing of DNA damage. As described above, *lem-3; brc-1* double mutants show increased sensitivity to IR compared to the single mutants (Fig. S1A). This loss of viability in the double mutant is associated with extensive chromatin bridge formation, and lagging chromosomes and micronuclei became apparent, especially upon real-time imaging (Fig. S4, Movie S4). It thus appears that LEM-3 is important to process DNA bridges that result from the incomplete processing of DNA breaks when BRC-1 function is compromised.

Other endonucleases such as MUS-81 and SLX-1-SLX-4 have been shown to resolve recombination intermediates during metaphase (for review^7^). To investigate the contribution of LEM-3 to recombination intermediates processing in relation to these other endonucleases, we assessed the progeny viability of *lem-3* and *slx-4* single and double mutants. SLX-4 scaffold protein interacts with MUS-81 and SLX-1 and provides a platform for recombination intermediate resolution. In *slx-4* mutants unresolved meiotic recombination intermediates are carried into the first zygotic division^21,22^. While *slx-4* single mutants still produce 20% viable embryos, *lem-3; slx-4* double mutants only produce inviable progeny (Fig. 3E). Imaging of the first zygotic cell division showed that chromatin bridges did occur in *slx-4* embryos, but were barely visible. They had no apparent effect on chromosome segregation based on distance measurements between separating nuclei, indicating that the chromatin bridges can be efficiently resolved (Fig. S5A and S5C). In contrast, chromosome segregation was severely compromised in *lem-3; slx-4* embryos, as indicated by the defective separation of nuclei and the formation of chromatin bridges and micronuclei (Fig. S5A, S5C and S5D). Interestingly, the separation of nuclei occurred faster in *lem-3* mutants with average nuclear separation rate of 5.31 μm/100s (standard deviation SD, ±0.13; n=5), compared to 4.02 μm/100s in wild type (SD, ±0.32; n=6, p<0.0001), 3.77 μm/100s in *slx-4* mutants (SD, ±0.23; n=6, p<0.00001), and 2.4 μm/100s in *lem-3; slx-4* double mutants (SD, ±0.37; n=5, p<0.00001), indicating a potential role for LEM-3 in delaying the separation of nuclei (Fig. S5A). Careful analysis revealed that in 32% of *lem-3; slx-4* embryos (n=19) cytokinesis failed, leading to binucleated cells and highly abnormal mitotic cell divisions (Fig. S5B and S5D). In summary, our data suggest that LEM-3 processes recombination intermediates, as well as DNA structures resulting from replication stress or partial chromosome decondensation, and that such processing helps to maintain genome integrity.

### The recruitment of LEM-3 to the midzone/midbody is important for chromatin bridge resolution

To better correlate the spatiotemporal behaviour of chromatin bridge resolution and the LEM-3 midbody localization, we simultaneously visualized the localization of YFP-LEM-3, the plasma membrane (mCherry-PH) and chromatin (mCherry-H2B) when chromatin bridges are induced by *capg-1* RNAi. We found that LEM-3 was first detectable at the midzone at the time of cleavage furrow completion, then increased in abundance and together with the chromatin bridges congressed into the midbody (Fig. S2B and S2C, white arrow; Movie S5). At that stage the bridges started to resolve and to retract, commencing from the midbody (Fig. S2B, white arrowheads, 810-830 seconds), consistent with the function of midbody tethered LEM-3.

To further test whether the ability of LEM-3 to resolve chromosome bridges requires an intact spindle midzone and the recruitment of LEM-3 to that site, we combined the partial *mcm-7* RNAi with the depletion of ZEN-4 and CYK-4 central spindle components. As expected we found that compromising central spindle formation abolished midzone localization of LEM-3 (Fig. 3F and S3B). Embryos solely treated with low dose of *mcm-7* RNAi never showed formation of chromatin bridges, as was the case for *zen-4* or *cyk-4* RNAi treated embryos (Fig. 3F and S3B). In contrast, when embryos treated with combination of partial *mcm-7* RNAi and *zen-4* or *cyk-4* RNAi, chromatin bridges formed with high penetrance in (4/7) and (4/9) cases, respectively (Fig. 3F and S3B, Movies S6 and S7). This finding indicates that the spindle midzone is important for the resolution of chromatin bridges, and by inference suggests that LEM-3 midbody localization might have a role in this.

### Regulation of LEM-3 midbody localization and function by AIR-2/Aurora B kinase

The Aurora B kinase dependent NoCut checkpoint is able to detect chromatin bridges and delay abscission, allowing time for chromatin bridge resolution^4,8,19^. We found that the *C. elegans* Aurora B homolog AIR-2 and LEM-3 partially co-localize at the midzone/midbody. Using Manders’ overlay coefficient 77.4%±9.3% of LEM-3 protein co-localized with AIR-2 at the midzone/midbody (Fig. 4A). In addition, LEM-3 localization depends on the AIR-2/Aurora B kinase (Fig. 4B), indicating that AIR-2 regulates the midbody association and/or activity of LEM-3. We thus searched for AIR-2/Aurora B consensus phosphorylation sites (K/R; K/R; X0-2; S/T) in the LEM-3 protein sequence and identified two serines, Ser192 and Ser194, embedded in such a consensus sequence (Fig. 4C). The corresponding sites are phosphorylated by AIR-2/Aurora B in ZEN-4/MKLP-4, and the first serine also occurs in human LEM-3 orthologue Ankle1^23^. We thus mutated serines 192 and 194 to alanine by genome engineering (Figure 5A). We observed comparable expression levels of GFP-LEM-3 S192AS194A and the wild type control (GFP-LEM-3, Fig. 5B). However, the *lem-3 S192AS194A* mutant showed an increased sensitivity to IR upon treatment of L4 stage larvae and late stage embryos (Fig. 5C and 5D). In addition, combining the *lem-3 S192AS194A* mutations with *slx-4* resulted in reduced viability compared to *slx-4* mutant, consistent with increased levels of chromatin bridges observed in *lem-3 S192AS194A; slx-4* double mutants (Fig. S6A and S6B). Careful examination of the localization of LEM-3 revealed no detectable LEM-3 foci at the midzone in the *GFP-LEM-3 S192AS194A* mutant during the first cell division (Fig. 5E, Movie S8). Upon condensin inactivation by *capg-1* RNAi, GFP-LEM-3 S192AS194A midbody location appeared later and reduced as compared to wild type (Fig. 5E, white arrows; Movie S9). Consistent with LEM-3 midbody location contributing to chromatin resolution, chromatin bridges persisted for longer when the LEM-3 phosphorylation mutant embryos were treated with *capg-1* RNAi (Fig. 5E, white arrowheads).

**Fig. 4.**
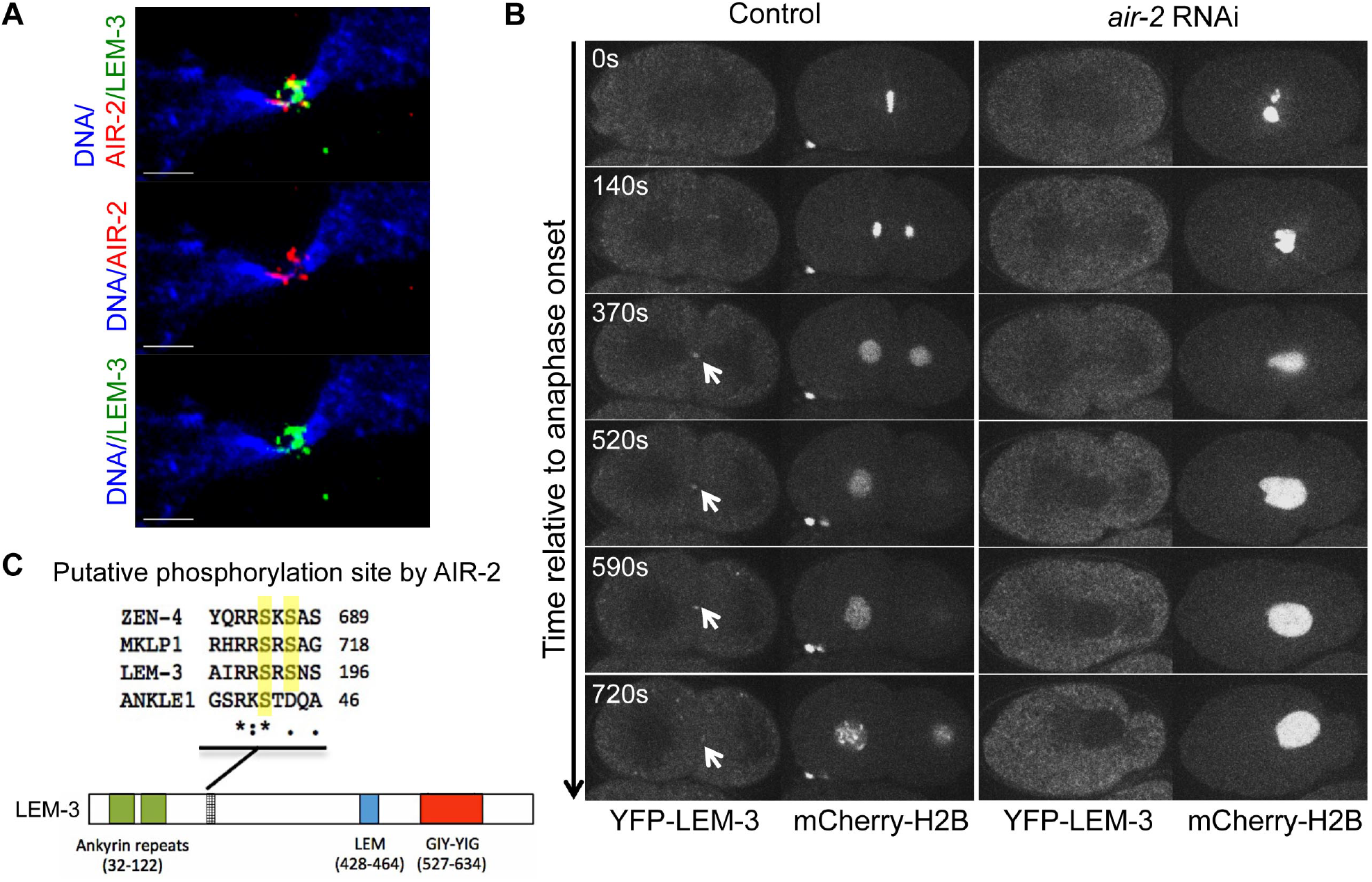
LEM-3 localization depends on the AIR-2/Aurora B kinase. (A) Partial co-localization of LEM-3 with AIR-2/Aurora B kinase. AIR-2 immunostaining of representative embryos expressing GFP-LEM-3. Scale bars: 1 μm. (B) LEM-3 foci cannot be detected in *air-2* RNAi embryos. Arrows indicate YFP-LEM-3. (C) Schematic showing of the position and conservation of the putative AIR-2/Aurora B kinase phosphorylation sites in central spindle component ZEN-4/MKLP1 and LEM-3/Ankle1. Conserved serines are highlighted in yellow.

**Fig. 5.**
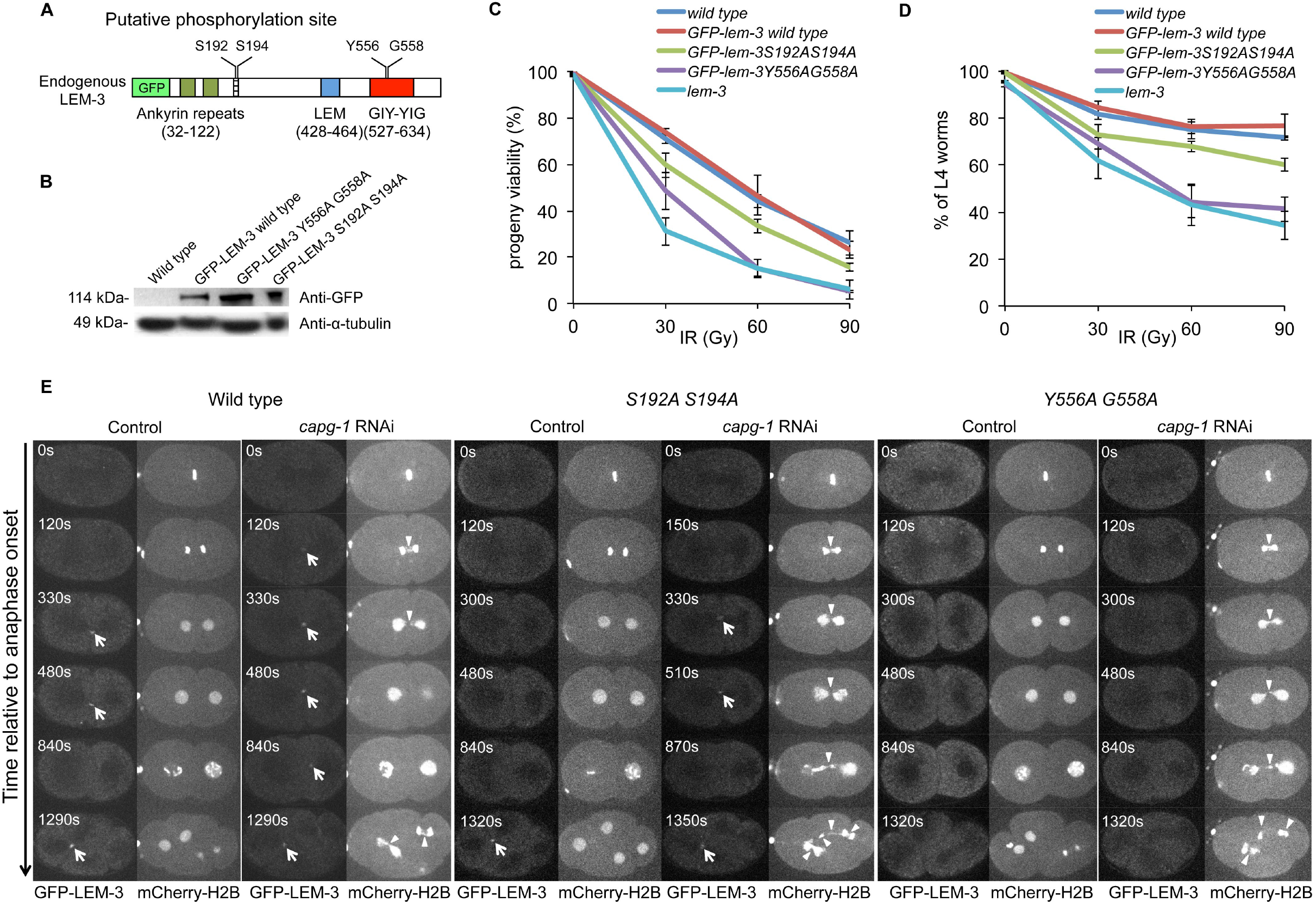
Functional dissection of LEM-3 phosphorylation sites and GIY-YIG motif. (A) Schematic representation of endogenous GFP-LEM-3 generated by CRISPR/Cas9-mediated genome editing. The conserved Serine 192 and 194 AIR-2 putative phosphorylation sites and Tyrosine 556 and Glycine 558 in the GIY-YIG motif are indicated. (B) Western blot analysis of GFP-LEM-3 protein levels in GFP-LEM-3 wild type, *GFP-lem-3 S192AS194A* and *GFP-lem-3 Y556AG558A* mutants. (C-D) Sensitivity of *GFP-lem-3 S192AS194A* and *GFP-lem-3 Y556AG558A* mutants to ionizing radiation (IR). Progeny viability of *lem-3* mutants (C) after IR treatment. (D) Development delay was scored 48 hours after IR by calculating the percentage of worms that reached the L4 stage. Values are the average of three independent experiments. Error bars represent standard deviation of the mean. (E) Analysis of chromosome segregation in *GFP-lem-3 S192AS194A* and *GFP-lem-3 Y556AG558A* mutants. Arrows indicate YFP-LEM-3. Arrowheads indicate the chromatin bridges.

To provide direct evidence that LEM-3 S192 and S194 are phosphorylated *in vivo*, we generated phospho-specific antibodies corresponding to a peptide containing both phosphorylated S192 and S194 (Fig. S7A). Using these LEM-3 phospho-specific antibodies we detected LEM-3 midbody-staining that overlapped with the localization of GFP-LEM-3 (Fig. 6A). The same co-localization occurred on chromatin bridges induced by *capg-1* and *mcm-7* RNAi (Fig. 6B and S7D). Importantly, no LEM-3 phosphorylation was detected on the midbody, when *lem-3 S192AS194A* and *lem-3 (tm3468)* mutants (the latter containing an inframe deletion that abrogates phosphorylation sites) were stained with the phospho-specific antibody (Fig. S7B and S7C). Whereas GFP-LEM-3 is detected between nuclei from anaphase onwards in the presence of chromatin bridges induced by *capg-1* RNAi, phosphorylated LEM-3 is only detected at the midbody during telophase (Fig. 6B). Therefore phosphorylation of LEM-3 might not be required for the initial recruitment but for congression of LEM-3 into the midbody. In summary, our data indicate that AIR-2 is required for robust localization of LEM-3, and LEM-3 phosphorylation may be important for stable association of LEM-3 with the midbody.

**Fig. 6.**
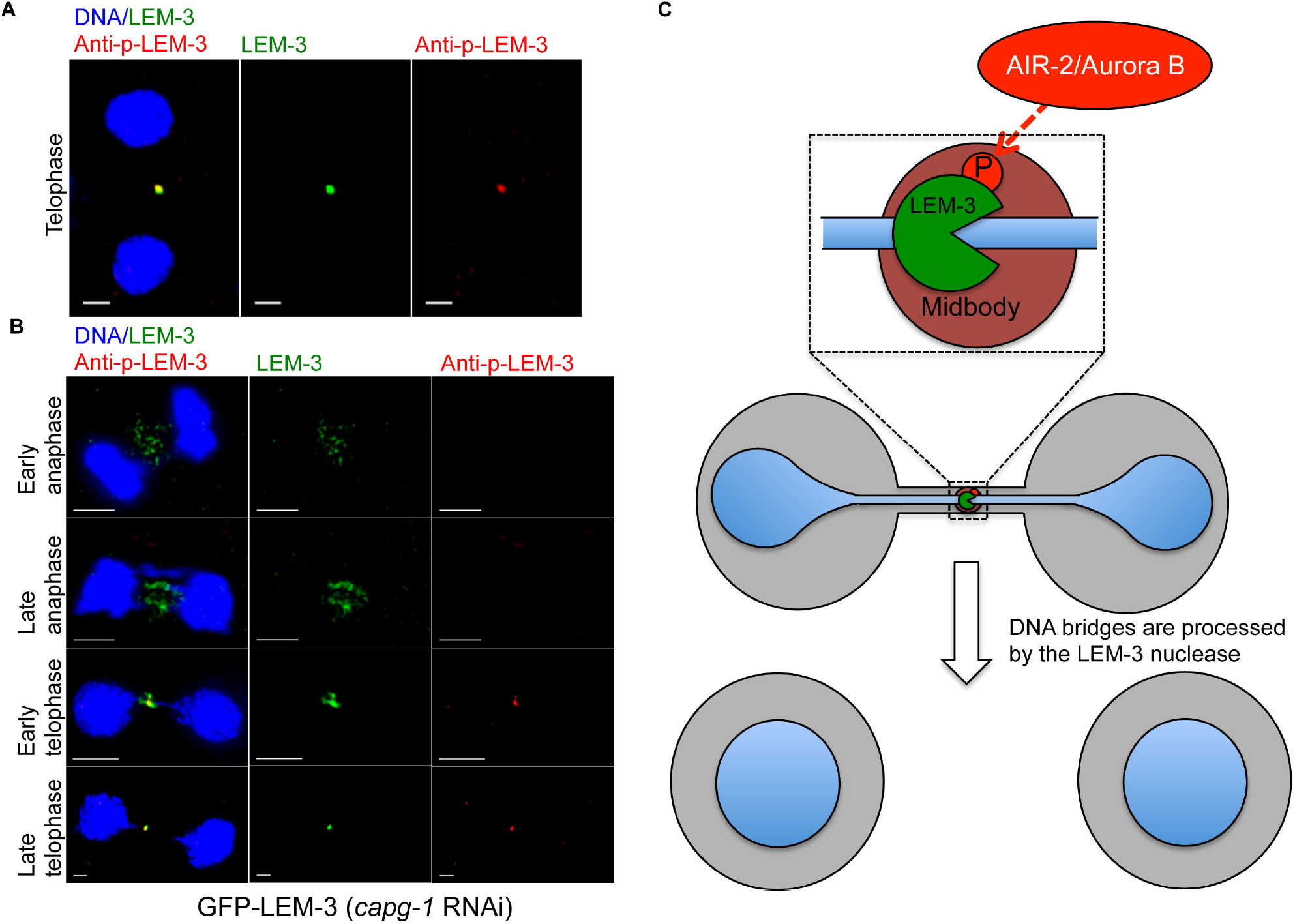
LEM-3 is phosphorylated at the midbody. (A) Co-localization of GFP-LEM-3 and phospho-LEM-3 at the midbody. Scale bars: 5 μm. (B) Localization of LEM-3 and phosphorylated LEM-3 in the presence of chromatin bridges from anaphase to telophase. (C) Proposed model for chromatin bridge processing by the midbody associated LEM-3 nuclease.

### The GIY-YIG nuclease motif is essential for *in vivo* function of LEM-3

To determine whether the conserved GIY-YIG nuclease motif is required for LEM-3 function, we mutated the catalytic Y556 and G558 residues, which are predicted to be essential for nuclease activity^24,25^. Interestingly, LEM-3 localization at the midzone/midbody was not detectable in the *Y556A G558A* double mutant even after induction of chromatin bridges by *capg-1* RNAi (Fig. 5E; S10) suggesting that the mutations we introduced into the GIY-YIG motif might also compromise DNA binding and or midbody localization. *lem-3 Y556A G558A* has the same level of protein expression as wild type but is hypersensitive to IR (Fig. 5B, 5C and 5D). Moreover, combining the *lem-3 Y556A G558A* mutation with *slx-4* led to 100% embryonic lethality and extensive chromatin bridge formation (Fig. S5A and S5B). In addition, excessive and persistent chromatin bridge formation occurred upon *capg-1* RNAi in *lem-3 Y556A G558A* embryos (Fig. 5E and Movie S10). These results suggest that the GIY-YIG motif is essential for LEM-3 function *in vivo*.

## Discussion

It has been known for a decade that the Aurora B mediated NoCut checkpoint plays a key role in delaying abscission, when chromatin is trapped at the midbody during cytokinesis^4,8^. However, it remained enigmatic how chromatin bridge resolution occurs and how this process is connected with the abscission machinery and the NoCut checkpoint. In this study we report that the conserved endonuclease LEM-3 accumulates at the midbody and provide evidence that it resolves chromatin bridges at the final stage of cell division (Fig. 6C). LEM-3 midbody localization requires central spindle formation and is regulated by AIR-2/Aurora B kinase (Fig. 6C). Together with our evidence that LEM-3 does not act within the known major DNA repair pathways, this suggests that LEM-3 defines a new mechanism for maintaining genome integrity that acts just before the final stage of cytokinesis to resolve chromatin bridges.

In addition to its role in delaying abscission in the presence of chromatin bridges by inducing a stable intercellular canal at the midbody, our data indicate that at the same time Aurora B also promotes bridge resolution via recruitment of LEM-3 to the midbody. We found that the worm Aurora B kinase AIR-2 co-localizes with LEM-3 and is required for LEM-3 association with the midbody (Fig. 4A and 4B). Although we have no evidence for the direct interaction, LEM-3 has a conserved AIR-2/Aurora B phosphorylation site and can be phosphorylated *in vivo*, suggesting that LEM-3 may be a substrate of AIR-2/Aurora B kinase. In addition, the occurrence of phosphorylated LEM-3 during late cytokinesis is consistent with the reported high level of AIR-2 kinase activity at this stage. It is noteworthy that the *lem-3 S192A S194A* phosphosite mutant is not as sensitive to IR as the *lem-3* null mutant. This might indicate that mutation of these two serines impairs but does not completely abolish LEM-3 function (Fig. 5C, 5D and 5E). This may be due to the residual midbody location of LEM-3 in this mutant (Fig. 5E). Our finding that midzone/midbody formation plays a key role in chromatin bridge resolution further supports the importance of LEM-3 midbody localization. The *C. elegans* checkpoint preferentially works after the second embryonic cell cycle^19^, rendering chromatin-bridge processing by LEM-3 more important in the first cell cycle. Indeed, the faster separation of nuclei we observed in *lem-3* mutants may suggest that LEM-3 also has a direct role in the NoCut checkpoint. Irrespectively, while we provide strong evidence that LEM-3 acts on the midbody during cytokinesis, we cannot rule out that LEM-3 might also have a role in earlier stages of the cell cycle.

We found that LEM-3 is able to bind to and resolve chromatin bridges caused by various perturbations, including incomplete DNA replication, unresolved recombination intermediates, and compromised chromosome condensation (Fig 1B, 3A, S2A and S5), indicating that LEM-3 can process a wide range of DNA substrates with distinct structures. In organisms with large genome, unreplicated DNA is frequently present in normal cells at the end of G2 phase^26^. The post-replicative resolution of unreplicated DNA depends on creation of ultrafine anaphase bridges during mitosis^26^. Ultrafine DNA bridges, which cannot be stained by conventional DNA dyes are different from chromatin bridges^27^. It is possible that the ultrafine DNA bridges exist in early *C. elegans* embryos where rapid cell divisions occur. Ultra fine bridges in mammalian cells are detected by being coated with the BLM or the PICH helicase^28-30^. We did not observe bridges coated with the *C. elegans* BLM ortholog HIM-6, and PICH has not been found encoded in the worm genome (our unpublished data). Irrespective, in order to segregate unreplicated segments of DNA, the unreplicated double stranded DNA has to be disentangled to form single stranded DNA, which can be repaired in the next cell cycle^26^. Therefore, the accumulation of LEM-3 at the midbody during a late stage of cell division may also contribute to the resolution of the ultrafine DNA bridges caused by unreplicated DNA regions in unperturbed cells.

In human cells, the cytoplasmic nuclease TREX1 was recently found to contribute to the cutting of chromatin bridges that originate from telomere fusion^31^. TREX1 binds to the chromatin bridges after anaphase, and its nuclease activity is important for their timely cleavage. Unlike LEM-3, whose localization becomes more restricted at the midbody as cytokinesis proceeds, TREX1 is present throughout chromatin bridges and generate extensive ssDNA. Moreover, TREX1 is an exonuclease that requires nicked DNA substrates, indicating that TREX1 is not solely responsible for chromatin bridge resolution, or else can only act on nicked DNA substrates. Furthermore, in contrast to LEM-3, which processes chromatin bridges at the end of cell division to maintain genome stability and allow for increased survival, this is not clear for TREX1, which might only get access to bridges when the nuclear envelope is perturbed. In addition, the cutting of telomere-telomere bridges by TREX1 leads to pathological ‘McClintock-like’ breakage-fusion-bridge cycles often linked with a large number of randomly assembled chromosome fragments, a phenomenon termed Chromothripsis^31–33^. In contrast, our data suggest that LEM-3 is able to process a variety of DNA bridges to promote viability (Fig. 3C and 3D). Thus LEM-3 is part of a ‘last chance saloon’ mechanism to maintain genome integrity.

It will be very interesting in future studies to explore further the role of the Ankle1 nuclease^10^, which is the ortholog of *C. elegans* LEM-3 in human and mouse. The localization of endogenous Ankle1 has neither been reported, nor its potential association with chromatin bridges explored, though over-expressed Ankle1 protein was found to be cytoplasmic^10^. Mice lacking Ankle1 do not have an overt phenotype and extracted cells are not hypersensitive to DNA damaging agents^11^, but this is also true of mice lacking the Gen1 or Mus81 nucleases, indicating a high level of redundancy between DNA nucleases that preserve genome integrity in mammalian systems^34^. Interestingly, increased susceptibility to breast and ovarian cancer in the general population and in carriers of BRCA1 and BRCA2 mutations was mapped down to 13 polymorphisms at 19p13.1 mapping to non-coding regions in the human LEM-3 homolog Ankle1 and a second gene ABHD8, encoding for a alpha beta hydrolase domain^13–15^. The effect of these polymorphisms on Ankle1 function remains to be explored, but our finding that *lem-3; brc-1* double mutants show elevated sensitivity to DNA damage (Fig. S1A and S4B) is consistent with Ankle1 being the most likely gene in 19p13.1 to affect breast cancer disposition. Indeed, another study found that coding mutations in Ankle1 were associated with increased risk of breast cancer^35^. It will be important in the future to explore whether Ankle1 and BRCA1 cooperatively promote genome stability in human cells, as predicted by the behaviour of their *C. elegans* equivalents.

## Methods

### Experimental Model and Subject Details

*C. elegans* strains were maintained at 20°C on nematode growth medium (NGM) plates seeded with OP50 bacteria. Transgenic lines were generated by injecting plasmid DNA/ssDNA directly into the hermaphrodite gonad. Integrated transgenic strains were outcrossed at least six times before use.

### Embryonic viability assay

L4 or young adult worms were treated with various DNA damaging agents as previously described^36^. After 24 hours of recovery, worms were transferred onto new plates and allowed to lay eggs for 6-8 hours. Eggs were quantified. Unhatched eggs were counted 24 hours later and the percentage of progeny viability was calculated.

### Embryo irradiation assay

The embryo irradiation assay was performed as described previously^17,36^. Experiments were done in triplicate. 10 young adult worms of each strain were transferred onto seeded plates. Worms were allowed to lay eggs for 2 h and were removed from the plate afterwards. The embryos were incubated at 20°C for 3 h before treatment with different doses of IR (0 Gy, 30 Gy, 60 Gy, 90 Gy). The eggs were counted, and the plates were incubated at 20°C for 24 h. Unhatched eggs were counted the next day and plates were incubated for another 24 h. The delayed development phenotype was scored 48 hours after IR by calculating the percentage of worms that reached the L4 stage.

### Image acquisition

For immunostaining, embryos were fixed in methanol at −20°C for 30 min. After three washes with PBST (0.1% Tween 20), samples were blocked with 0.1% BSA, incubated with primary antibodies overnight, and washed three times. Following a two-hour incubation with secondary antibodies and/or DAPI, cover glasses were rinsed five times before image analysis. Primary and secondary antibodies were used at the indicated dilutions: rabbit anti-AIR-2 (1:200)^37^, sheep anti-phospho-LEM-3 (1:100), Alexa 568 labelled donkey anti-rabbit (1:750), and Alexa 594 labelled donkey anti-sheep (1:1000) (Invitrogen). For super resolution OMX microscopy, established protocols were followed^38^. To record mitotic divisions, we followed previously described imaging procedures^18^. Embryos were dissected in M9 (3 g/l KH2PO4, 6 g/l Na2HPO4, 5 g/l NaCl, 1 mM MgSO4) and mounted on 2% agarose pads. Images were captured every 10 s at 23–24°C using a spinning-disk confocal microscope (IX81; Olympus) with spinning-disk head (CSU-X1; Yokogawa Electric Corporation) and MetaMorph software (Molecular Devices). Image analysis and video processing were performed using ImageJ software (National Institutes of Health).

#### RNAi

Clones for *mcm-7, capg-1, top-2, nmy-2, spd-1, zen-4* and *cyk-4* feeding strains were obtained from Source BioScience. To make RNAi plates, bacteria were grown to 0D600 0.6-0.8. After induction with 1 mM IPTG for 4 hours at 37°C, bacteria were spread on NGM plates. After 16 hours L4 worms were transferred to RNAi-plates and fed for at least 24 hours at 25°C before analysis. For double RNAi, equal amounts of bacterial cultures were mixed prior to seeding the RNAi plates. Partial *mcm-7* and *capg-1* RNAi were achieved by diluting *mcm-7* or *capg-1* RNAi expressing bacteria with control bacteria^18^.

### Generation of point mutations by CRISPR/Cas9 genome editing

The *cop859 [Plem-3::eGFP::STag:: lem-3::3’UTRlem-3*] eGFP insertion was generated by Knudra (<u>http://www.knudra.com/</u>) following the procedures described by Dickinson et al.^*39*^. Exact details are available upon request. For generated point mutations in the endogenous *lem-3* locus, we performed CRISPR-Cas9 following procedures described by the Seydoux lab^40^ using modified protocol provided by Simone Köhler and Abby Dernburg. The injection mixture was prepared by pre-assembling gRNA-Cas9 ribonucleoprotein (RNP) complexes *in vitro*. 1:1 mixture of tracrRNA and crRNA (100 μM each) were heated at 95 °C for 5 min, followed by annealing at room temperature for 5 min. Cas9-RNP complex was then formed by adding purified Cas9 protein (IDT) into a injection mixture (10 μl) with a final concentration of 30 μM gRNA and 28 μM Cas9 protein and incubated at room temperature for 20 min. Repair templates were provided as 100 ng/μl ssDNA. 2.5 ng/μl of pCFJ90 and 5 ng/μl of pCFJ104 plasmids were used as red fluorescent co-injection makers. 10-40 young adult hermaphrodites were injected for each mutant and recovered onto individual plate. Fluorescent F1 worms were picked after 2-3 days post-injection and screened by PCR or DNA sequencing. crRNA sequence for *GFP-lem-3S192A194A* was 5’-AAGAAGTAGATCCAACAGCA-3’, and the repair template sequence (ssDNA, Ultramer oligo, IDT) was as follows: cgttcaatcgaggaaatgttcttactaattcatatagatgtgcaaagaagaaaatcagagccacatttcatgctattcgaagaGCtagaGcTaacagcacCgcaacacttcaagacgttgttttaacatctgaaggaattcgaacagtgaccacacctagtaggagagcacctaaggcaaccgtttatgc.

crRNA sequence for *GFP-lem-3 Y556AG558A* was 5’-GTACGATCAATCTTTTACGT-3’, and the repair template sequence (ssDNA, Ultramer oligo, IDT) was ttcggatataatgcgttttgctatctcattatggatcctcgaattttgggaagcaatgtggagaaccttacacttgaaacctttgtacgatcGatctttGCcgtagCTaaaggttcaaaaaatcgtccattagctcatttcattgatgctcgcaatgaacggagagacaaattggataaactaaaaacttgcgaaaag.

### Western blotting

Wild type, *GFP-lem-3, GFP-lem-3 S192AS194A* and *GFP-lem-3 Y556AY558A* adult worms were collected and bleached until they were completely lysed. After centrifugation for 1 min at 2200 rpm the pellet was resuspended in 200 μL M9 worm buffer. The suspension was transferred into safety capped 1.5 mL reaction tubes. 200 μL of 40% TCA and 300 μL of glass beads were added and the worm eggs were disrupted by bead beating for 2 min. The suspension was then transferred into a 1.5 mL reaction tube. The beads were washed with 5% TCA which was added to the suspension afterwards. Samples were spun down for 10 min at 3000 rpm, the supernatant was removed and the pellet was resuspended in 1 mL ice-cold acetone. After another centrifugation for 2 min at 11,000 rpm the acetone was discarded and the pellet was left to dry at room temperature. Finally the pellet was resuspended in 100 μL Laemmli buffer (3x). 10 μL of each sample was loaded onto a 4%-12% NuPage precast gel. For Western blotting primary and secondary antibodies were used at the indicated dilutions: mouse anti-GFP (1:1000), mouse anti-α-tubulin (1:5000), horse anti-mouse IgG-HRP (1:3000) (NEB).

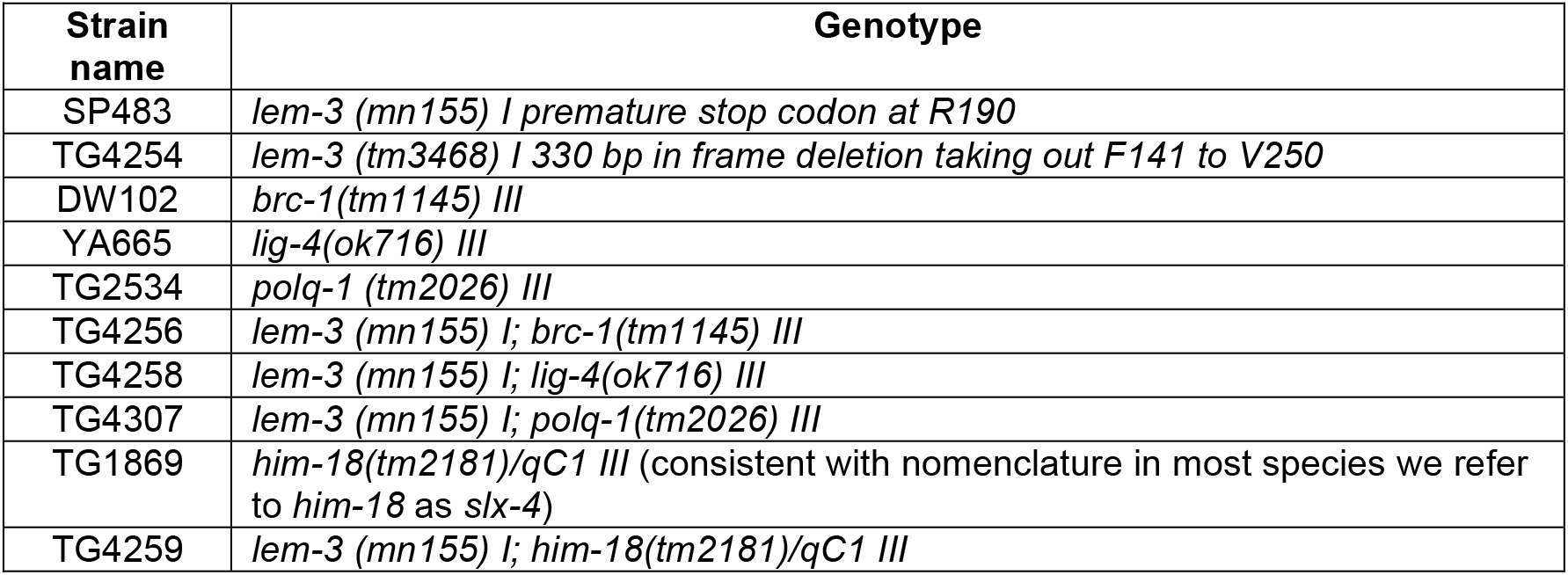

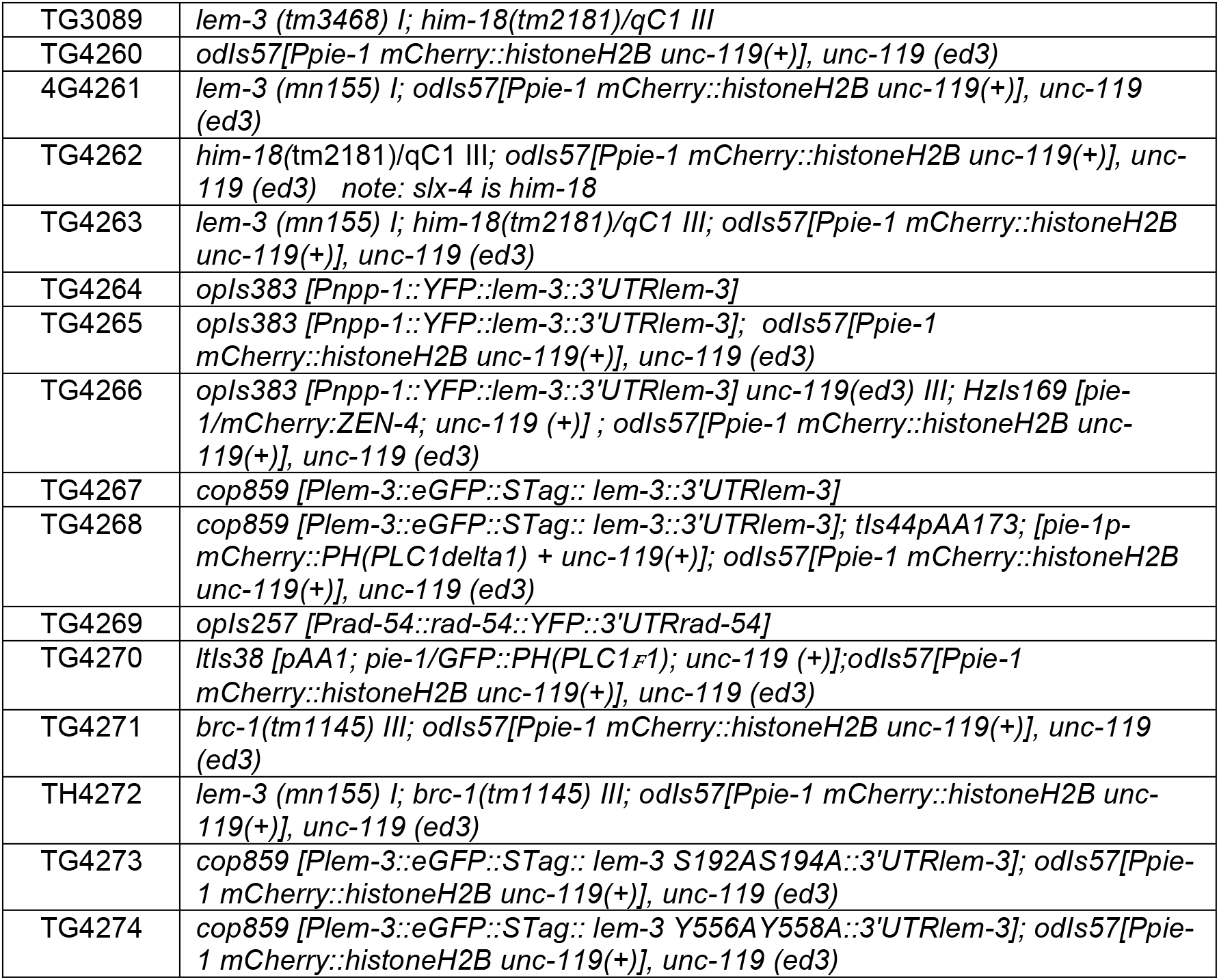
List of strains

## Acknowledgements

This work was funded by Wellcome Trust Programme (AG 0909444/Z/09/Z), Investigator (KL 102943/Z/13/Z) and Strategic awards (097045/B/11/Z), a MRC core grant KL (MC_UU_12016/13) and the FWF (VJ SFB-F34). VS and YH were supported by a Wellcome PhD fellowship and ISSF funds. The Dundee Imaging Facility was supported by the Wellcome Trust (097945/B/11/Z) and the MRC (MR/K015869/1). We thank the Caenorhabditis Genetics Center (funded by NIH Office of Research Infrastructure Programs P40 OD010440) for worm strains. We thank Federico Pelisch for the AIR-2 antibody, James Hastie and Fiona Brown for the phospho-specific LEM-3 antibody (MRC PPU reagents, https://mrcppureagents.dundee.ac.uk). We are grateful to Graeme Ball, Callum Tromans-Coia and Markus Posch for technical assistance. We thank Simone Köhler and Abby Dernburg for sharing genome-engineering protocols. We thank Ulrike Gartner for proofreading. Experiments were mostly done by YH, with contributions from RS BW VS BM AW. YH, AG, VJ and KL wrote the paper.

---

**Fig. S1.**
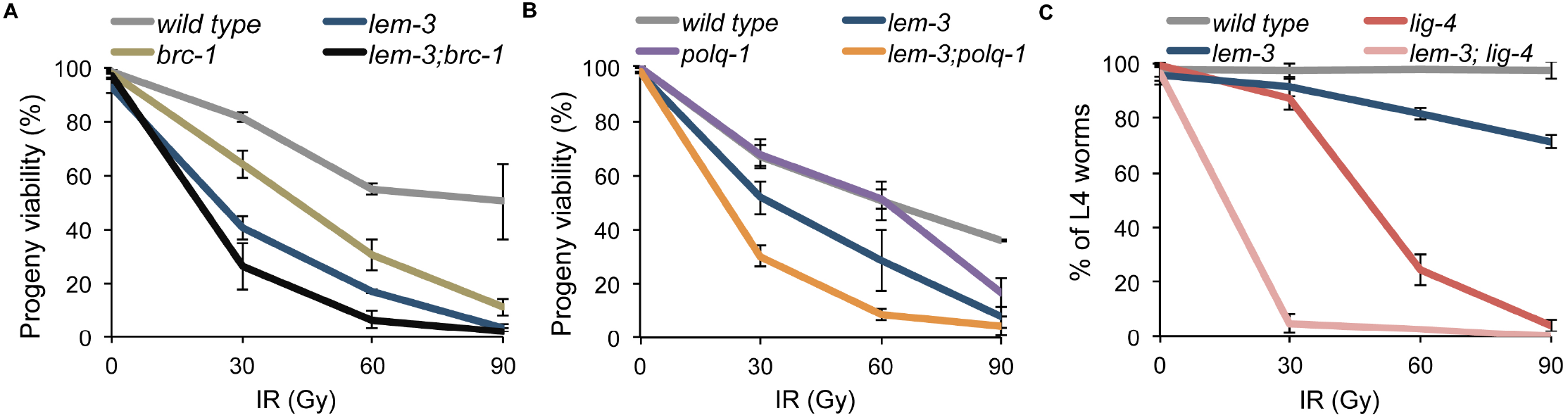
Sensitivity of *lem-3* single and double mutants to IR. (A and B) Young adult worms were treated with ionizing radiation. After 24 hours of recovery, worms were transferred onto new plates and allowed to lay eggs for 6-8 hours. Progeny viability in % was determined by counting number of viable eggs/total number of eggs laid. (C) Late stage embryos were irradiated and the proportion of animals researching L4 stage was scored. This assay accounts for defect in end joining a pathway that predominantly acts in *C. elegans* somatic tissues. Error bars represent standard deviation of the mean.

**Fig. S2.**
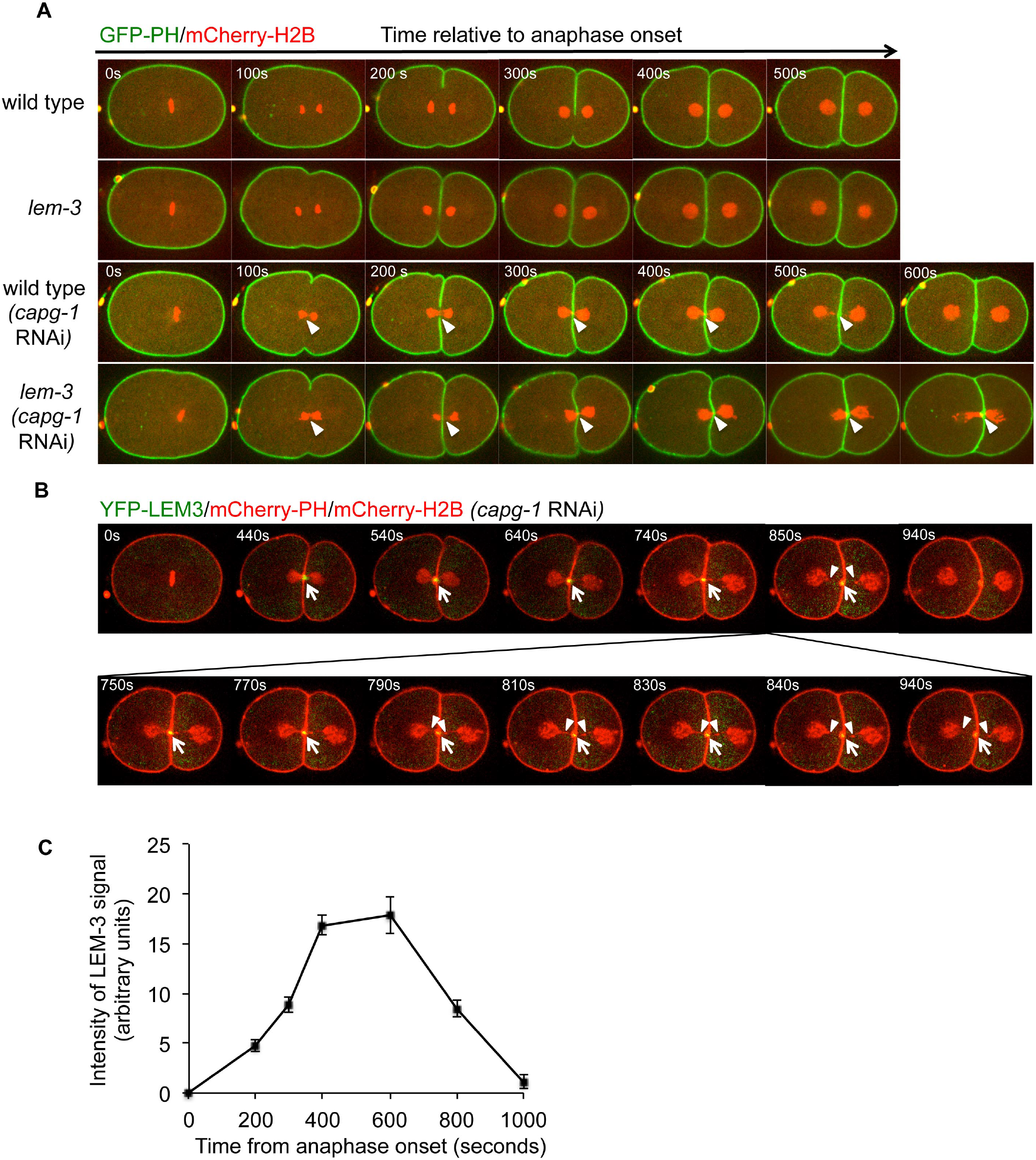
Persistent *capg-1* RNAi induced chromatin bridges in *lem-3* mutants. (A) Persistent *capg-1* RNAi induced chromatin bridges in *lem-3* mutants. Chromosome segregation in wild type and *lem-3* mutant embryos upon depletion of the condensin I subunit CAPG-1 (100% *capg-1* RNAi). Images were taken from control and *capg-1* RNAi embryos expressing mCherry-H2B and GFP-PH. Arrowheads indicate chromatin bridges. Times are relative to anaphase onset of the first division. (B) Localization of LEM-3 before and after resolution of chromatin bridges induced by *capg-1* RNAi. Arrows indicate YFP-LEM-3. Arrowheads indicate retracting chromatin bridges. (C) Quantification of intensity of LEM-3 signal at the midbody during chromatin bridge resolution.

**Fig. S3.**
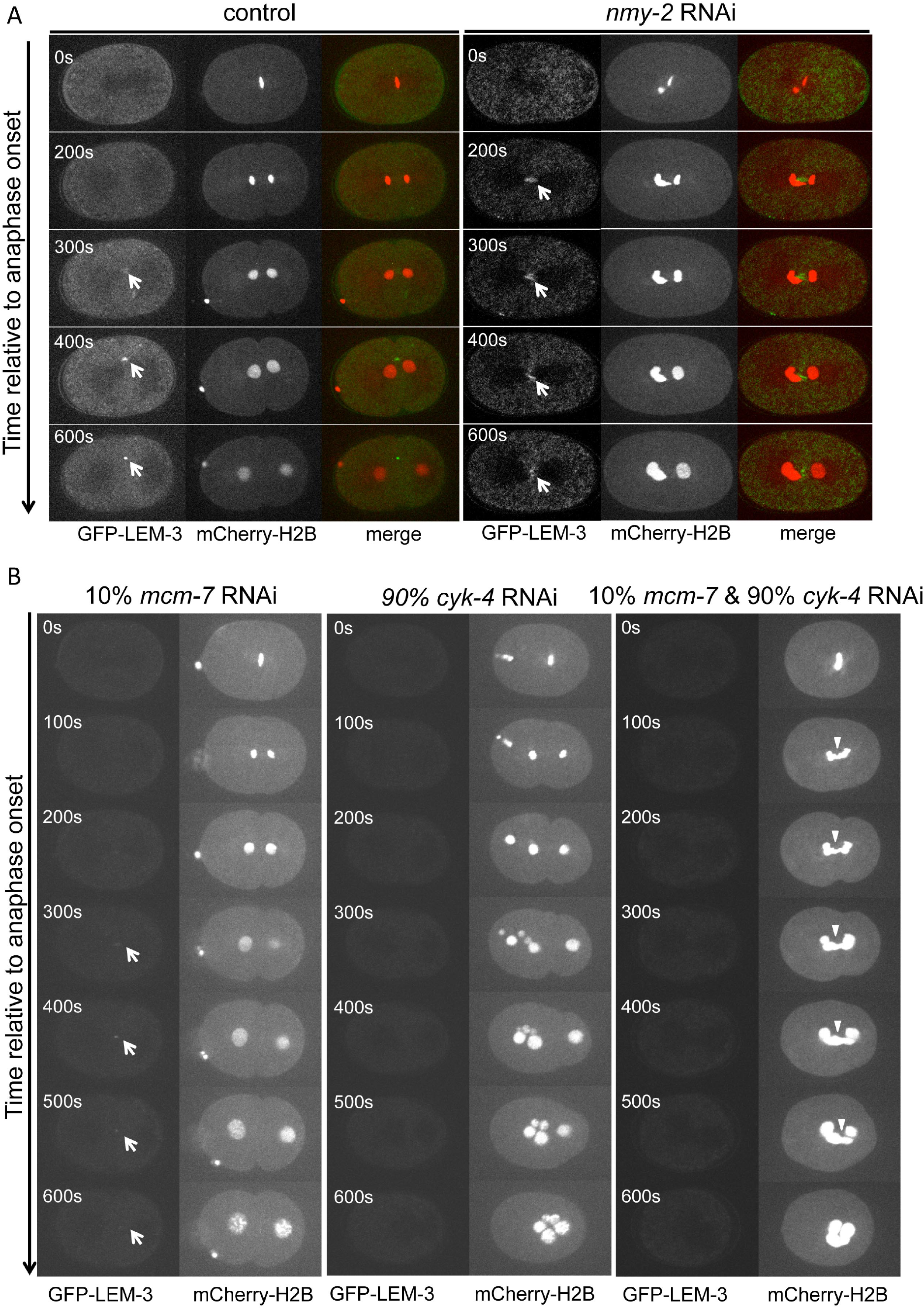
The recruitment of LEM-3 to the midbody is important for chromatin bridge resolution. (A) Midbody localization of LEM-3 is independent on the contractile ring formation. Depletion of the contractile ring component NMY-2 by *nmy-2* RNAi didn’t affect the recruitment of LEM-3 to the midbody. Images were taken from a time-lapse recording of embryos expressing GFP-LEM-3 and mCherry-H2B from the anaphase onset of the first mitotic division. Times are relative to anaphase onset. (B) Formation of chromatin bridges upon partial depletion of DNA replication helicase subunit MCM-7 and central spindle component CYK-4. Arrows indicate GFP-LEM-3. Arrowheads indicate chromatin bridges.

**Fig. S4.**
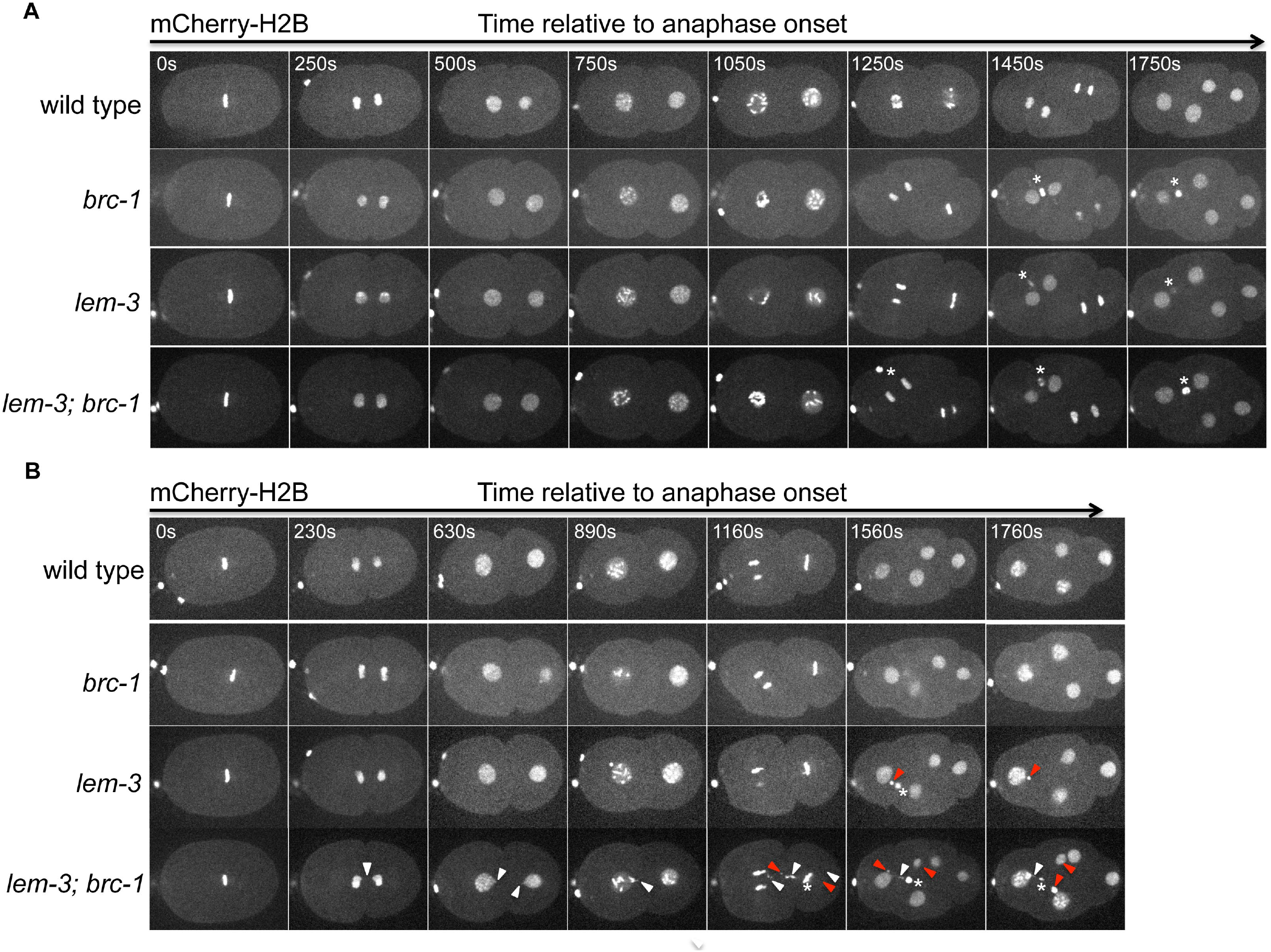
LEM-3 and BRC-1 are required for faithful chromosome segregation after IR. (A) Worms of the indicated genotypes were dissected and one cell stage embryos were used for recording. Images were taken from time-lapse recordings of embryos expressing mCherry-H2B. Times are relative to anaphase onset of the first division. (B) L4 stage worms were irradiated with 30 Gy. After 24 hours adult worms were dissected and embryos were used for recording. White arrowheads indicate chromatin bridges, red arrowheads indicate micronuclei. Stars indicate polar bodies.

**Fig. S5.**
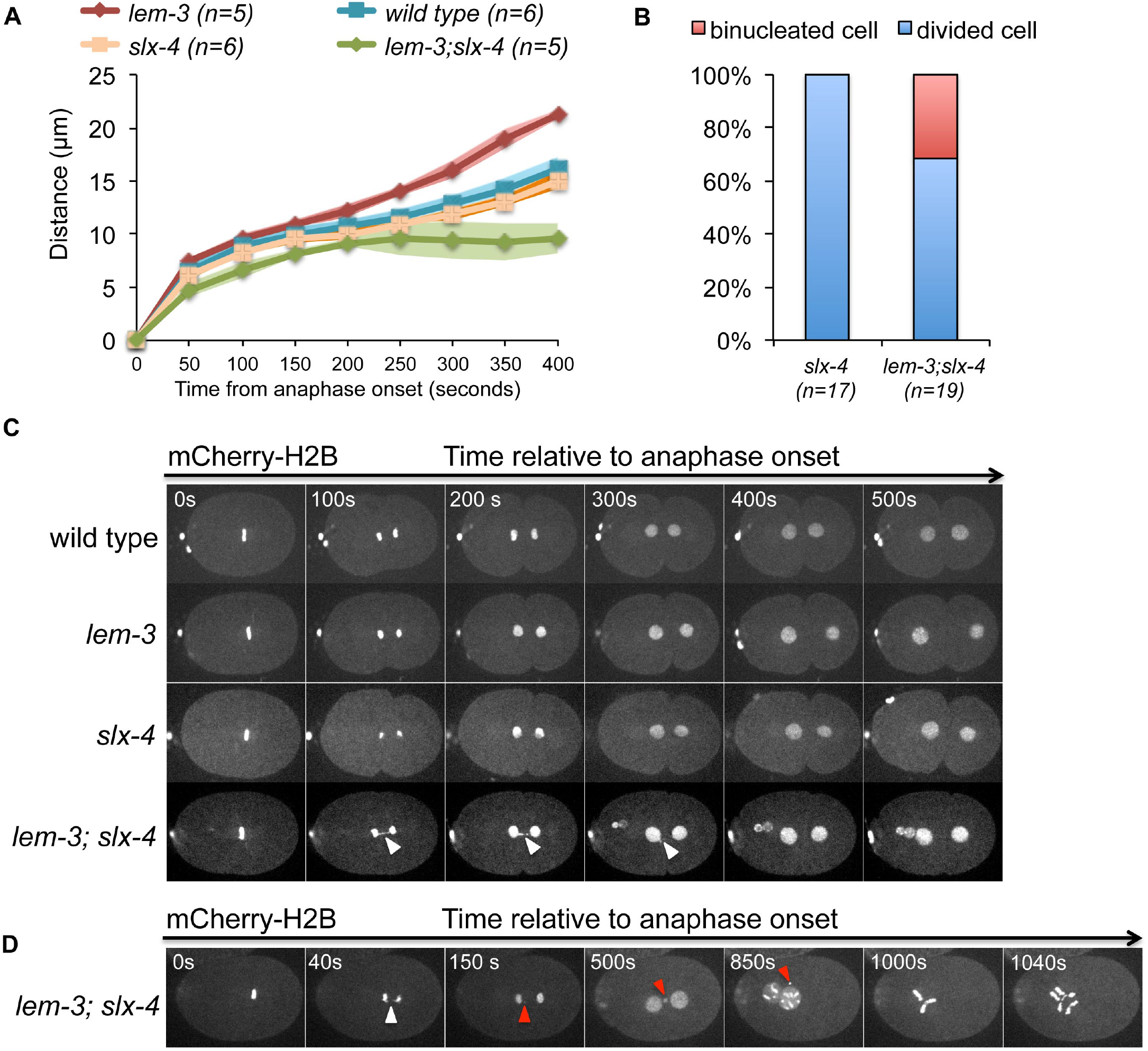
Chromosome segregation defects in *lem-3; slx-4* double mutants. (A) Analysis of chromosome segregation in wild type, *lem-3, slx-4* single mutants and *lem-3; slx-4* double mutants by measuring the distance between two separated nuclei. The sample size (n) indicates the total number of embryos examined for each genotype. (B) Quantification of binucleated cell formation in *lem-3; slx-4* double mutants. The sample size (n) indicates the total number of embryos examined for each genotype. (C) Chromosome segregation in embryos of indicated genotypes. Representative images were taken from time-lapse recordings of an embryo expressing mCherry-H2B from the anaphase onset of first mitotic division. White arrowheads indicate chromatin bridges. (D) Example of binucleated cells observed in *lem-3; slx-4* double mutants. Images were taken from a time-lapse recording of an embryo expressing mCherry-H2B from the anaphase onset of first mitotic division. White arrowheads indicate chromatin bridges. Red arrowheads indicate micronuclei. The formation of a tripolar spindle is evident ~1000 seconds after the first anaphase.

**Fig. S6.**
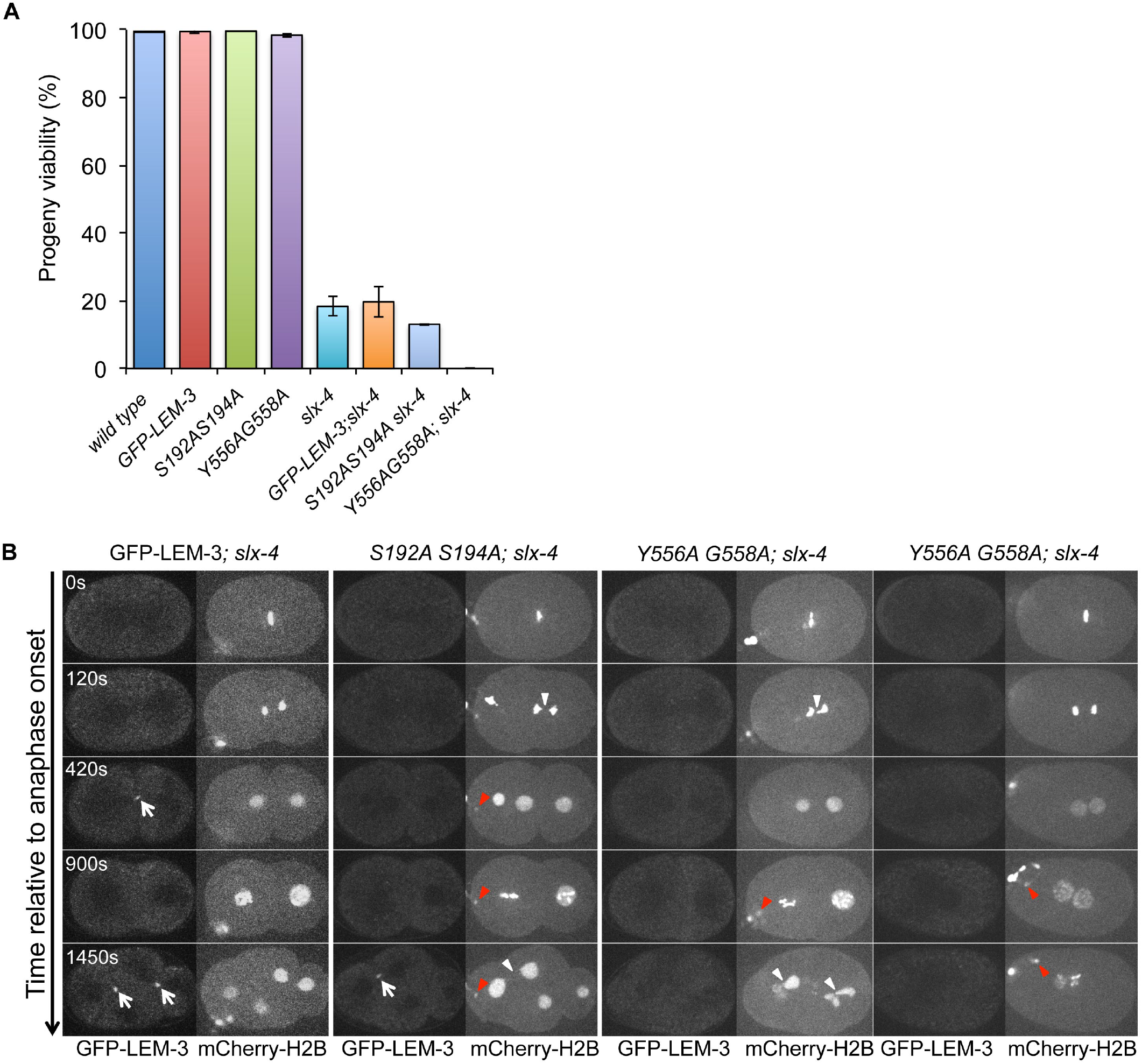
Compromised chromosome segregation in *GFP-lem-3 S192AS194A; slx-4* and *GFP-lem-3 Y556AG558A; slx-4* double mutants. (A) Progeny viability of animals of the indicated genotypes. Percentage survival progeny was determined by counting number of viable eggs/total number of eggs laid. Error bars represent standard deviation of the mean. (B) Chromosome segregation in *GFP-lem-3 S192AS194A; slx-4* and *GFP-lem-3 Y556AG558A; slx-4* double mutants. Images were taken from time-lapse recordings of embryos expressing mCherry-H2B from the anaphase onset of first mitotic division. Arrows indicate GFP-LEM-3. White arrowheads indicate chromatin bridges. Red arrowheads indicate micronuclei.

**Fig. S7.**
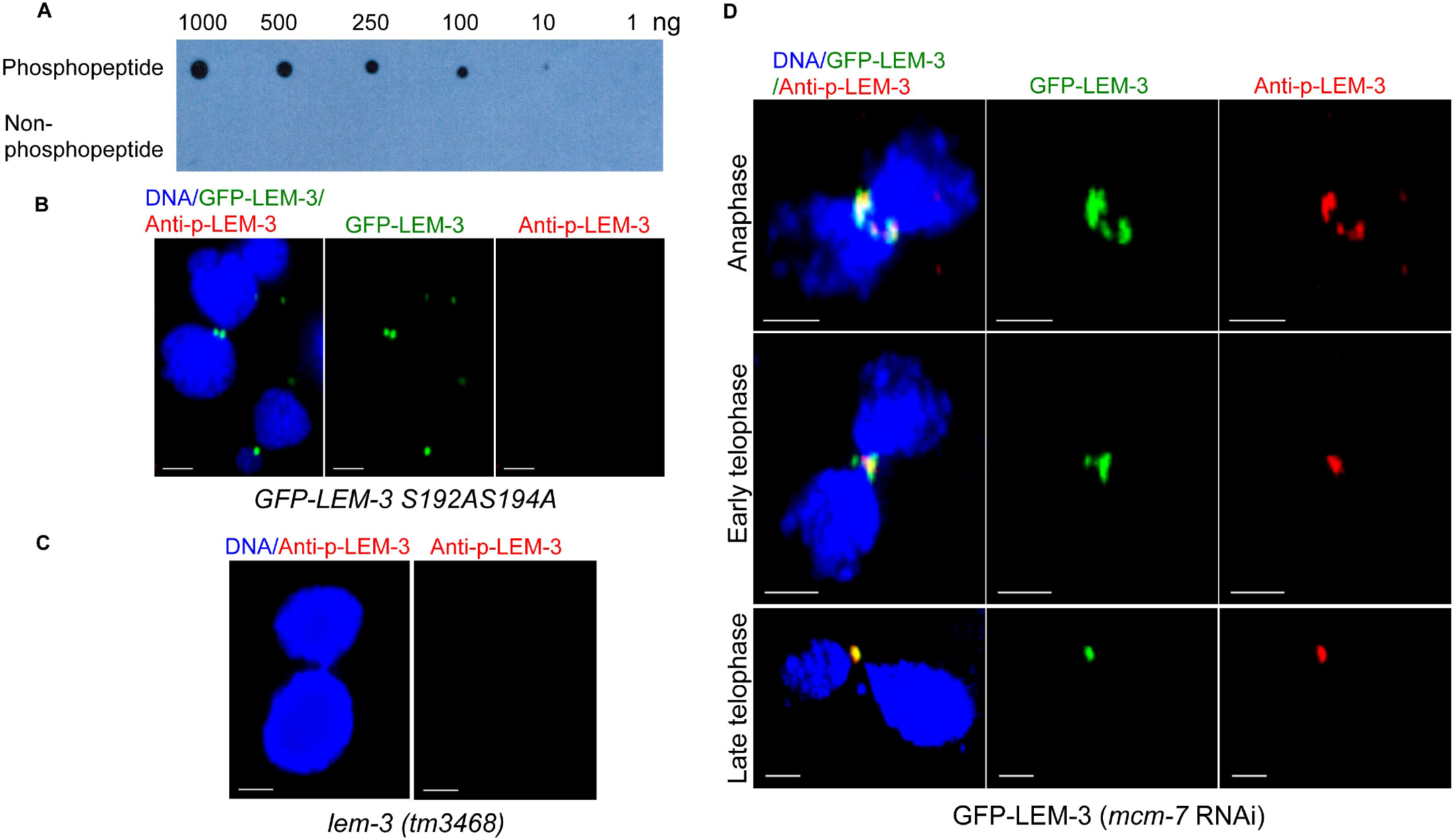
Specificity analysis of phospho-LEM-3 antibody. (A) Dot blot analysis to test specificity towards LEM-3 phospho-peptides and non-phospho-peptides. An increasing amount of the phospho- and non-phospho-peptide was spotted on nitrocellulose and probed with a 1:500 dilution of the anti-phospho LEM-3 antibody and a ten-fold dilution of non-phospho-peptide as blocking agent. (B) Immunostaining of phospho-LEM-3 in *GFP-lem-3 S192AS194A* mutant embryos. (C) Immunostaining of phospho-LEM-3 in *lem-3 (tm3468)* embryos. (D) Representative images of GFP-LEM-3 co-localizing with phospho-LEM-3 at midzone /midbody in the presence of chromatin bridges induced by *mcm-7* RNAi. Scale bars: 2 *μ* m.

**Movie S1. Localization of YFP-LEM-3 during cell division**

**Movie S2. Co-localization of YFP-LEM-3 and mCherry-ZEN-4 at the midbody**

**Movie S3. YFP-LEM-3 localization upon *mcm-7* RNAi**

**Movie S4. Aberrant chromosome segregation in *lem-3; brc-1* mutants upon treatment with IR**

**Movie S5. Chromatin bridge resolution by LEM-3**

**Movie S6. Chromatin bridge formation upon partial *mcm-7* and *zen-4* RNAi.**

**Movie S7. Chromatin bridge formation upon partial *mcm-7* and *cyk-4* RNAi.**

**Movie S8. Localization of GFP-LEM-3 S192A S194A**

**Movie S9. GFP-LEM-3 S192A S194A localization upon *capg-1* RNAi**

**Movie S10. GFP-LEM-3 Y556A G558A localization upon *capg-1* RNAi**

## References

1 Shi, Q. & King, R. W. Chromosome nondisjunction yields tetraploid rather than aneuploid cells in human cell lines. Nature 437, 1038–1042, (2005).

2 Fujiwara, T. et al. Cytokinesis failure generating tetraploids promotes tumorigenesis in p53-null cells. Nature 437, 1043–1047, (2005).

3 Hoffelder, D. R. et al. Resolution of anaphase bridges in cancer cells. Chromosoma 112, 389–397, (2004).

4 Steigemann, P. et al. Aurora B-mediated abscission checkpoint protects against tetraploidization. Cell 136, 473–484, (2009).

5 Liu, Y., Nielsen, C. F., Yao, Q. & Hickson, I. D. The origins and processing of ultra fine anaphase DNA bridges. Curr. Opin. Genet. Dev. 26, 1–5, (2014).

6 Chan, K. L., North, P. S. & Hickson, I. D. BLM is required for faithful chromosome segregation and its localization defines a class of ultrafine anaphase bridges. EMBO J 26, 3397–3409, (2007).

7 Matos, J. & West, S. C. Holliday junction resolution: regulation in space and time. DNA Repair 19, 176–181, (2014).

8 Norden, C. et al. The NoCut pathway links completion of cytokinesis to spindle midzone function to prevent chromosome breakage. Cell 125, 85–98, (2006).

9 Dittrich, C. M. et al. LEM-3 - A LEM domain containing nuclease involved in the DNA damage response in C. elegans. PLoS One 7, e24555, (2012).

10 Brachner, A. et al. The endonuclease Ankle1 requires its LEM and GIY-YIG motifs for DNA cleavage in vivo. J. Cell Sci 125, 1048–1057, (2012).

11 Braun, J., Meixner, A., Brachner, A. & Foisner, R. The GIY-YIG type endonuclease ankyrin repeat and LEM domain-containing protein 1 (ANKLE1) is dispensable for mouse hematopoiesis. PLoS One 11, e0152278, (2016).

12 Zlopasa, L., Brachner, A. & Foisner, R. Nucleo-cytoplasmic shuttling of the endonuclease ankyrin repeats and LEM domain-containing protein 1 (Ankle1) is mediated by canonical nuclear export- and nuclear import signals. BMC Cell Biol 17, 23, (2016).

13 Antoniou, A. C. et al. A locus on 19p13 modifies risk of breast cancer in BRCA1 mutation carriers and is associated with hormone receptor-negative breast cancer in the general population. Nature Genet. 42, 885–892, (2010).

14 Bolton, K. L. et al. Common variants at 19p13 are associated with susceptibility to ovarian cancer. Nature Genet. 42, 880–884, (2010).

15 Lawrenson, K. et al. Functional mechanisms underlying pleiotropic risk alleles at the 19p13.1 breast-ovarian cancer susceptibility locus. Nature Commum. 7, 12675, (2016).

16 Lemmens, B. B. & Tijsterman, M. DNA double-strand break repair in *Caenorhabditis elegans*. Chromosoma 120, 1–21, (2011).

17 Clejan, I., Boerckel, J. & Ahmed, S. Developmental modulation of nonhomologous end joining in *Caenorhabditis elegans*. Genetics 173, 1301–1317, (2006).

18 Sonneville, R., Craig, G., Labib, K., Gartner, A. & Blow, J. J. Both chromosome decondensation and condensation are dependent on DNA replication in *C. elegans* embryos. Cell Rep. 12, 405–417, (2015).

19 Bembenek, J. N., Verbrugghe, K. J., Khanikar, J., Csankovszki, G. & Chan, R. C. Condensin and the spindle midzone prevent cytokinesis failure induced by chromatin bridges in *C. elegans* embryos. Curr. Biol. 23, 937–946, (2013).

20 Pavicic-Kaltenbrunner, V., Mishima, M. & Glotzer, M. Cooperative assembly of CYK-4/MgcRacGAP and ZEN-4/MKLP1 to form the centralspindlin complex. Mol. Biol..Cell 18, 4992–5003, (2007).

21 Agostinho, A. et al. Combinatorial regulation of meiotic holliday junction resolution in *C. elegans* by HIM-6 (BLM) helicase, SLX-4, and the SLX-1, MUS-81 and XPF-1 nucleases. PLoS Genet. 9, e1003591, (2013).

22 Saito, T. T., Lui, D. Y., Kim, H. M., Meyer, K. & Colaiacovo, M. P. Interplay between structure-specific endonucleases for crossover control during Caenorhabditis elegans meiosis. PLoS Genet 9, e1003586, (2013).

23 Guse, A., Mishima, M. & Glotzer, M. Phosphorylation of ZEN-4/MKLP1 by aurora B regulates completion of cytokinesis. Curr. Biol. 15, 778–786, (2005).

24 Kowalski, J. C. et al. Configuration of the catalytic GIY-YIG domain of intron endonuclease I-TevI: coincidence of computational and molecular findings. Nucleic Acids Res. 27, 2115–2125 (1999).

25 Truglio, J. J. et al. Structural insights into the first incision reaction during nucleotide excision repair. EMBO J. 24, 885–894, (2005).

26 Moreno, A. et al. Unreplicated DNA remaining from unperturbed S phases passes through mitosis for resolution in daughter cells. Proc. Natl. Acad. Sci.USA 113, 5757–5764, (2016).

27 Mankouri, H. W., Huttner, D. & Hickson, I. D. How unfinished business from S-phase affects mitosis and beyond. EMBO J. 32, 2661–2671, doi:10.1038/emboj.2013.211 (2013).

28 Biebricher, A. et al. PICH: a DNA translocase specially adapted for processing anaphase bridge DNA. Mol. Cell 51, 691–701, (2013).

29 Chan, K. L. & Hickson, I. D. New insights into the formation and resolution of ultra-fine anaphase bridges. Semin. Cell Dev Biol. 22, 906–912, (2011).

30 Ke, Y. et al. PICH and BLM limit histone association with anaphase centromeric DNA threads and promote their resolution. EMBO J. 30, 3309–3321, (2011).

31 Maciejowski, J., Li, Y., Bosco, N., Campbell, P. J. & de Lange, T. Chromothripsis and kataegis induced by telomere Crisis. Cell 163, 1641–1654, (2015).

32 Li, Y. et al. Constitutional and somatic rearrangement of chromosome 21 in acute lymphoblastic leukaemia. Nature 508, 98–102, (2014).

33 Meier, B. et al. C. elegans whole-genome sequencing reveals mutational signatures related to carcinogens and DNA repair deficiency. Genome Res. 24, 1624–1636, (2014).

34 Wang, X. et al. Gen1 and Eme1 play redundant roles in DNA repair and meiotic recombination in mice. DNA Cell Biol. 35, 585–590,

35 Rhie, S. K. et al. Comprehensive functional annotation of seventy-one breast cancer risk Loci. PLoS One 8, e63925, (2013).

36 Craig, A. L., Moser, S. C., Bailly, A. P. & Gartner, A. Methods for studying the DNA damage response in the *Caenorhabdatis elegans* germ line. Methods Cell Biol. 107, 321–352, (2012).

37 Pelisch, F. et al. A SUMO-dependent protein network regulates chromosome congression during oocyte meiosis. Mol. cell 65, 66–77, (2017).

38 Hong, Y. et al. The SMC-5/6 Complex and the HIM-6 (BLM) helicase synergistically promote meiotic recombination intermediate processing and chromosome maturation during *Caenorhabditis elegans* meiosis. PLoS Genet. 12, (2016).

39 Dickinson, D. J., Pani, A. M., Heppert, J. K., Higgins, C. D. & Goldstein, B. Streamlined genome engineering with a self-excising drug selection cassette. Genetics 200, 1035–1049, (2015).

40 Paix, A., Folkmann, A., Rasoloson, D. & Seydoux, G. High Efficiency, Homology-directed genome editing in *Caenorhabditis elegans* using CRISPR-Cas9 ribonucleoprotein complexes. Genetics 201, 47–54, (2015).

